# Liposomal DAGLβ Inhibitor Shows Substantial Antinociceptive Potency Mediated by AMPK in a Neuropathic Pain Mouse Models

**DOI:** 10.64898/2026.10.15.758837

**Authors:** Liam A. Babcock, Madeline Gunawardena, Payton E. Lowrey, Rudra Pangeni, McKenzie Hall, Michael Taylor, Saniya Chethan, Sara M. Herz, Dayna A. Adeso, Olivia L. Murtagh, Qingguo Xu, Ku-Lung Hsu, Aron H. Lichtman

**Affiliations:** Department of Pharmacology & Toxicology, Virginia Commonwealth University, Richmond, VA, USA, 23298, USA; Department of Pharmaceutics, Virginia Commonwealth University, Richmond, VA, 23298, USA; Department of Chemistry, University of Texas at Austin, Austin, TX, 78712, USA; Department of Pharmaceutical Sciences, Philadelphia College of Osteopathic Medicine, Suwanee, GA, 30024, USA; Department of Ophthalmology, Biomedical Engineering, Pediatrics, Center for Drug Discovery, and Massey Comprehensive Cancer Center, Virginia Commonwealth University, Richmond, VA, 23298, USA

**Keywords:** KT109, Antinociception, Cannabinoid, AMPK, Inflammation, Analgesic

## Abstract

Diacylglycerol lipase-beta (DAGLβ), a biosynthetic enzyme for the endocannabinoid 2-arachidonoylglycerol (2-AG), is highly expressed in macrophages and serves as a potential contributor to inflammatory and hypernociceptive responses. Moreover, the DAGLβ inhibitor KT109 produces antinociceptive effects in various rodent pain models, and its liposome formulation dramatically increases its antinociceptive potency in a lipopolysaccharide mouse model of inflammatory pain compared to free KT109. Here, we further optimized the liposomal KT109 formulation through adjustment of lipid composition and improved processing, achieving 100-fold higher drug encapsulation efficiency than the previous formulation, uniform size distribution, and excellent storage stability. Further, the second generation liposomal KT109 showed remarkable potency (ED50 [95% CL] = 0.226 [0.113-0.454] mg/kg) in a mouse model of paclitaxel-induced neuropathic pain, compared to free KT109 (ED50 [95% CL] = 31 [21–46] mg/kg). Liposomal KT109 (0.4 mg/kg) fully reversed paclitaxel-induced mechanical hypersensitivity for up to 24 hours, and its antinociceptive effects were sustained following a week of daily injections, with no detectable tolerance. In addition, the adenosine monophosphate-activated protein kinase (AMPK) inhibitor, dorsomorphin, prevented the antinociceptive effects of KT109, which is consistent with previous work demonstrating AMPK activation mediates the antinociceptive phenotype of DAGLβ knockout mice.

**Graphical Abstract:** 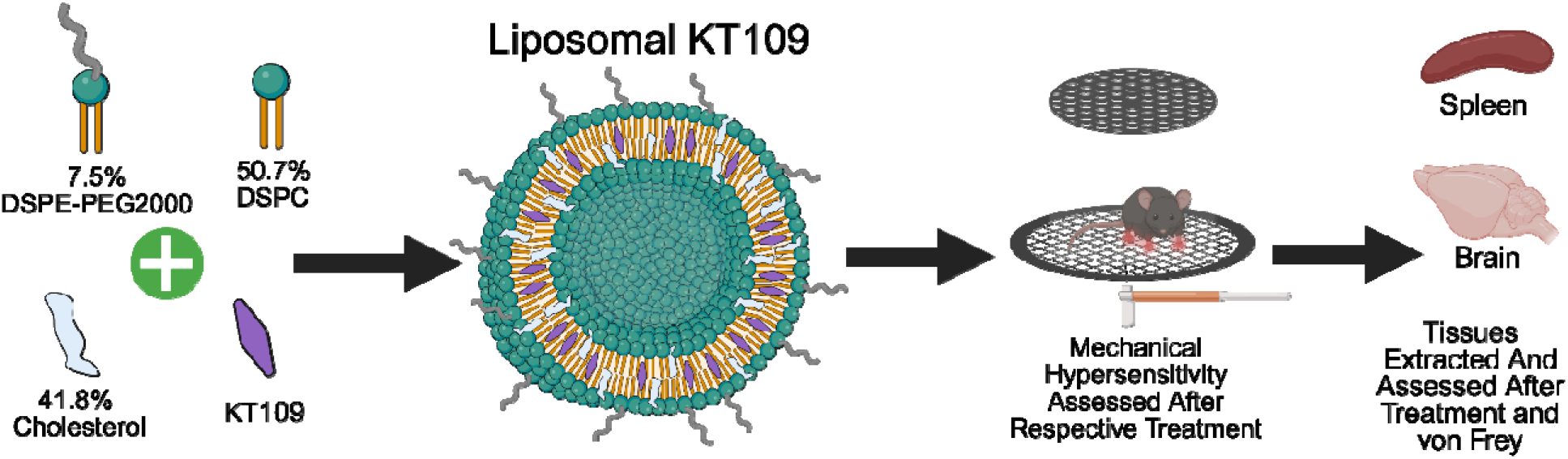

## 1. Introduction

Cancer remains a leading global health crisis, with an estimated 2,114,850 new diagnoses and 626,140 deaths projected to occur in the United States alone in 2026 [1]. Despite the significant advancements that have been made with cancer treatments over the past several decades, commonly used chemotherapeutics (e.g. vinca alkaloids, taxanes, platinum compounds, thalidomide, and bortezomib) often induce a phenomenon known as chemotherapy-induced peripheral neuropathy (CIPN). Symptoms of CIPN include tingling sensations, numbness, and pain, with these sensations being particularly concentrated in the hands and feet for some patients [2–4]. Moreover, these symptoms can persist for years after completing treatment, leading to chronic pain and a series of other debilitating health conditions [3,4]. Currently, treatment options for CIPN-associated pain remain limited, with common treatments (e.g., duloxetine) only providing modest efficacy. Other options to treat CIPN could be limiting the chemotherapy dose or using a different chemotherapeutic. Thus, there is a need to identify novel therapeutic strategies for the management of CIPN-associated pain. Here we examine whether inhibition of diacylglycerol lipase beta (DAGLβ), a biosynthetic enzyme for the endocannabinoid 2-arachidonylglycerol (2-AG) represents a viable strategy to ameliorate hypernociceptive responses in an established mouse model of CIPN.

Considerable work has demonstrated that the DAGLβ-selective inhibitor, KT109 [5], or genetic deletion of DAGLβ produces antinociceptive effects in CIPN, sickle cell disease, lipopolysaccharide (LPS) induced allodynia, and sciatic nerve injury rodent models of pain [6,7]. Despite the antinociceptive efficacy of KT109 in multiple models of inflammatory and neuropathic pain, its hydrophobic nature makes clinically relevant formulations difficult to produce. Given the high expression of DAGLβ in macrophages and considering that liposomes are inherently taken up by mononuclear phagocytes, our team developed a liposomal KT109 formulation made up of DSPC, DSPE-PEG2000, and DOPE with a KT109 encapsulation efficiency (EE) of 0.4%, and demonstrated that it had significantly increased *in vivo* antinociceptive potency compared to free KT109 delivered in a standard vehicle in an LPS mouse model of inflammatory pain [8]. Although this first generation liposomal KT109 formulation demonstrated the potential of the liposomal delivery system to improve delivery of KT109, the composition used led to poor formulation stability and low drug EE, underscoring the need for further optimization to support better translational feasibility [8]. Thus, our goal was to incorporate KT109 into a more clinically relevant, FDA-approved lipid composition [9,10], characterize its *in vitro* and *in situ* performance, and ultimately, to assess its *in vivo* efficacy for treating neuropathic pain using a CIPN mouse model.

Here, we first improved the liposomal KT109 formulation by adjusting the lipid composition [9–11] and processing. We then compared the size distribution and storage stability of this formulation to the first-generation formulation. After developing an improved liposomal KT109 formulation, we confirmed that this formulation, similar to previously published liposomal KT109, operated as a DAGLβ inhibitor in basal and LPS-activated macrophages [8]. We further expand upon this investigation by comparing free and liposomal KT109 inhibition *in situ* to understand the influence that liposomal delivery had on previously established off-target reactivity of KT109 [5].

Following inhibitory confirmation, we began our *in vivo* investigation of liposomal KT109 in CIPN utilizing the von Frey test. It is well-established that the hypersensitivity to non-noxious stimuli observed through the von Frey test is a hallmark of the dysregulated nociception seen in many pain models including CIPN [6]. Since the previous liposomal KT109 formulation greatly enhanced *in vivo* efficacy in LPS pain models [8], we hypothesized that our improved liposomal KT109 formulation would also greatly out-perform free KT109 in the CIPN model. We further utilized this method to elucidate the duration and efficacy with repeated administration to expand our understanding of translational feasibility. We used competitive activity-based protein profiling (ABPP) to confirm minimal off-target sites of the second generation liposomal KT109, which has been demonstrated with the previous generation of liposomal KT109.

Previous work has shown that dorsomorphin, an adenosine monophosphate-activated protein kinase (AMPK) inhibitor, reversed the antinociceptive phenotype of DAGLβ knockout (KO) mice in the LPS inflammatory model of pain suggesting that AMPK activation plays an essential role in the antinociceptive effects produced by DAGLβ inhibition [12]. Therefore, with our improved liposomal KT109 formulation, we anticipated that AMPK activation would also play an essential role in the antinociceptive effects produced. We tested whether AMPK activation mediates the observed antinociception of liposomal KT109 through pre-treatment with dorsomorphin. In this study, we developed a new and improved liposomal formulation of the DAGLβ inhibitor, KT109. To evaluate the suitability of this formulation as a future treatment for CIPN, we formulated liposomes with optimized composition for KT109 encapsulation, studied their delivery of KT109 in macrophages and tissues, and assessed the therapeutic effect of this formulation using the von Frey test in a CIPN mouse model.

## 2. Materials and Methods

### 2.1 Materials

All Lipids used for liposomal preparation were purchased from reputable suppliers. 1,2-Distearoyl-sn-glycero-3-phosphocholine (DSPC; COATSOME MC-8080; >99% purity) and N-(Methylpolyoxyethylene oxycarbonyl)-1,2-distearoyl-sn-glycero-3-phosphoethanolamine, sodium salt (DSPE-PEG2000; SUNBRIGHT DSPE-020CN; >99% purity) were purchased from NOF American Corporation. Cholesterol (≥ 92.5% purity) was purchased from Sigma-Aldrich.

### 2.2 Preparation of Liposomal KT109

All batches of liposomal KT109 were prepared using the thin-film hydration method. To begin, each of the lipids (400 mg DSPC, 210.6 mg DSPE-PEG2000, and 161.2 mg Cholesterol) were weighed and dissolved in 40 mL 1:1 v/v mixture of chloroform:methanol. This solvent composition was selected based on drug and lipid solubility. 10 mL of stock solution containing 1 mg/mL KT109, dissolved in the same solvent system, was then added to the lipid solution before addition to a 250 mL round-bottom glass flask. Solvent evaporation was then performed using a Buchi Rotovapor R-100 system, resulting in creation of the thin film. The film was allowed to dry overnight to ensure complete solvent evaporation, then rehydrated with 20 mL of PBS under vortex and extruded through filter membranes, ranging from 800nm to 100nm size using the Genizer GJE-10mL Jacketed Liposome Extruder. To ensure complete conversion from multi-lamellar lipid vesicles (MLVs) to nanoscale large lamellar vesicles (LUVs), the formulation was passed through each membrane size five times. Finally, the liposomal suspensions were added to Amicon Ultra Centrifugal Filters (10kDa, regenerated cellulose, 15 mL sample volume) and washed three times (4°C, 5000 RPM for 30 mins each, with fresh PBS added to replace volume lost after each wash) to remove excess material. Liposomal blank formulation directly mirrored the liposomal KT109 formulation procedure, apart from adding the drug solution. All liposomal solutions were stored at 4°C prior to use.

### 2.3 In Vitro Characterization of Liposomal KT109

#### 2.3.1 Evaluation of Physicochemical Properties and Morphology

Physicochemical properties of the liposomal formulations, including average droplet size, polydispersity index (PDI), and zeta potential, were evaluated using standard dynamic light scattering methods (Malvern Zetasizer Nano ZS90; Malvern Instruments, Malvern, UK) by diluting the liposomal suspensions in 10 mM NaCl solution (1:100 v/v dilution). Liposomal concentration was quantified for KT109 loaded formulations using nanoparticle tracking analysis (NTA) with a ZetaView PMX130 MONO (Particle Metrix, Germany) as previously described in previous study detailing liposomal blank concentration [11]. Morphologies of the liposomal suspensions were assessed using cryo-TEM as previously described [11,13]. Briefly, the formulations were applied to freshly glow discharged 200 mesh copper grids with a Lacey Carbon Film and a continuous layer of Ultrathin Carbon Film (Electron Microscopy Sciences, SKU:LC200-Cu-CC), blotted for 7 s with Whatman 1 Filter Papers at 100% relative humidity and 20 °C and plunged into liquid nitrogen-cooled ethane using an FEI Vitrobot MarkIV (Thermo Fisher Scientific). Images were collected on a Thermo Fisher Scientific Glacios cryo-TEM (Thermo Fischer Scientific) equipped with a Falcon 4i detector with a pixel size of 1.91 Å, and a total exposure time of 21 s resulting in a total accumulated dose of 30 e-/Å2 at a rate of 1.44 e-/Å2/s. The cryo-TEM data processing was performed in cryoSPARC v4.7.1 [14].

#### 2.3.2 Encapsulation Efficiency and Drug Content

KT109 content within the liposomal samples was quantified using high-performance liquid chromatography methods (HPLC-UV, Shimadzu Prominence LC System), equipped with an Agilent Pursuit XRs 5 C18 column (250 x 4.6 mm) with UV-detector system. The liposomal suspension was dissolved in 1:1 v/v acetonitrile:methanol at 1:500 v/v ratio, then the solution was filtered through 0.22 µm membrane filters (PTFE, hydrophobic, ThermoScientific) and 10 µL solution was injected into the HPLC. Mobile phases consisted of acetonitrile and water at a ratio of 90:10 v/v, both containing 0.1% trifluoroacetic acid, and the flow rate was set at 1.2 mL/min with detection at 279 nm. This procedure was used to quantify drug loading, as well as to evaluate drug content within the samples at each stability timepoint.

#### 2.3.3 Storage Stability Assessment of Liposomal KT109

Stability studies for the liposomal KT109 and blank formulations were performed in accordance with ICH Q1A(R2) and FDA recommendations for liposomal drug products [15]. Formulations were stored as aqueous suspensions at refrigerated conditions (5 ± 3°C), protected from light, and sampled at predetermined intervals (1, 7, 30, 45, 60, 90, and 120 days). At each time point, the main critical quality attributes - including particle size, PDI, zeta potential, drug content, and visual appearance - were analyzed to assess physical and chemical stability. All relevant attributes were characterized using previously described methods.

### 2.4 In Situ Evaluation with Liposomal KT109

#### 2.4.1 Cell Culture

RAW264.7 cells were cultured at 37°C under 5% CO_2_ in RPMI-1640 medium containing 10% fetal bovine serum (FBS, US Source, Omega Scientific), 1% L-glutamine (Fisher Scientific), and 100 µg/mL penicillin-streptomycin (Sigma-Aldrich). Following *in situ* vehicle (PBS), Liposomal KT109 (200 nM), or KT109 (200 nM) treatments on confluent cells for 24hr and LPS (5µg/mL) or PBS treatment (2hr), cells were harvested, washed three times in cold DPBS (400 x g, 3 min), and stored in -80°C.

#### 2.4.2 Preparation of Cellular and Tissue Proteomes

Directly following von Frey threshold assessment and euthanasia, relevant tissues were immediately extracted, weighed, and snap frozen. Cells were resuspended in small volumes of PBS. RAW264.7 cells were lysed via probe sonication (20% amplitude; Fisher Scientific Sonic Dismembrator FB120). Both brain and spleen tissues were rinsed in petri dish containing PBS and dried on filter paper wherein a 20-30mg portion was cut, weighed in a tared microcentrifuge tube. Tissues were dounce homogenized on ice in 1mL douncers (Kimble) at 50 mg/mL for brains and spleens, centrifuged at 100 x g for 5 min to remove cellular debris, and the supernatant was collected. In all cases, resulting lysate was fractionated via centrifugation at 100,000 x g for 45 min at 4 °C. Insoluble and soluble fractions were separated. Insoluble fractions were resuspended in PBS. Protein concentrations for each fraction were determined using a Bio-Rad DC protein assay.

#### 2.4.3 Quantitative LC-MS/MS Chemoproteomic Sample Preparation and Analysis

1 mg of proteome from each sample was treated with FP-Biotin for 2 hr (10 µM) at room temperature. Samples were then prepared as previously described. Peptides were acquired with nano-electrospray ionization-liquid chromatography-mass spectrometry (LC-MS/MS) on a Vanquish Neo UHPLC coupled to an Orbitrap Eclipse Tribrid mass spectrometer as previously described [11,16]. Data including Log_2_(Fold Change) and p-values are provided in **Supplemental Data Set 1.**

### 2.5 Evaluation of Efficacy in Chemotherapy-Induced Peripheral Neuropathy

#### 2.5.1 Animals

Adult male and female C57BL/6J mice (Jackson Laboratory, Bay Harbor, ME, USA) were used in all experiments. Mice of the same sex were group housed four per cage in a temperature (20–22 °C), humidity (55 ± 10%), and light-controlled (12-hour light/dark; lights on at 0600) AAALAC-approved (Protocol#: AD10000369) vivarium at Virginia Commonwealth University (VCU), with standard rodent chow and water available ad libitum. Animal studies were conducted in compliance with the National Institutes of Health Guide for the Care and Use of Laboratory Animals and approved by the VCU Institutional Animal Care and Use Committee. All animal interactions, including behavioral experimentation, occurred between the hours of 8am and 6pm. At each study endpoint, mice were either euthanized via cervical dislocation or CO2 asphyxia, followed promptly by cervical dislocation.

#### 2.5.2 Animal Handling and Acclimation

For all behavioral experiments, mice were acclimated to our vivarium with minimal disturbance upon arrival for one week prior to handling. Following vivarium acclimation, each mouse was handled in the vivarium for 2 minutes per day for four consecutive days, with animal scruffing performed every other day starting on day 4, which continued until experimentation began. Starting on Day 5, mice were acclimated daily to the testing space for 4 hours with handling in the testing room in a manner consistent with vivarium handling until experimentation started. During von Frey wire mesh acclimation days, handling would be performed after wire mesh acclimation. Frequent handling continued throughout all experimentation in an equal manner in the testing space and the vivarium.

#### 2.5.3 Behavioral Assessment of Mechanical Hypersensitivity

Mechanical hypersensitivity was assessed using the von Frey assay. Mice were placed unrestrained in individual inverted ventilated Plexiglas cylinders (approximately 8 cm in diameter) on a wire mesh screen (spaces 0.5 mm apart) and covered with a sheet that blocks light to allow better acclimation to the test environment. For all experiments, mice were habituated to the von Frey wire mesh screen as described above for approximately 2 hours per day for three consecutive days prior to experimentation. To determine paw withdrawal thresholds, calibrated von Frey monofilaments varying in mechanical stimulus intensity were applied randomly to both hind paws three times successively using a modified up and down stepwise method described as follows. The von Frey monofilaments in grams of force (gf) tested are 0.07, 0.4, 0.6, 1, 1.4, 2, and 4 gf. Any instance of lifting, licking, or shaking the paw was defined as a response. The stepwise von Frey test began with application of the 0.6 gf filament to both hind paws. For paws that showed a positive response to 0.6 gf, the force was decreased to 0.16 gf. If the response remained positive, the force was further decreased to a minimum of 0.07 gf. If the response to 0.16 gf was negative, the force was increased to 0.4 gf which is the final lower threshold tested. If there was no response to the initial 0.6 gf stimulus, the force was increased to 1.4 gf. A positive response to 1.4 gf prompted testing with the 1 gf filament, whereas a negative response resulted in an increase to 2 gf. If no response was observed at 2 gf, the force was progressively increased up to a maximum of 4 gf which was the final higher threshold tested. Paw withdrawal thresholds from both hind paws were averaged for each mouse. On test days, mice were acclimated to a wire mesh screen in individual Plexiglass cylinders for 1 hour, assessed for baseline thresholds, and then given their respective treatment via intraperitoneal injection.

#### 2.5.4 Drug Preparation for Animal Injections

Paclitaxel was dissolved in a 1:1:18 (v/v/v) solution (5% 200 proof ethanol, 5% emulphor, and 90% of 0.9% saline) to achieve a concentration of 0.8 mg/ml. KT109 and dorsomorphin free drug formulations were prepared by suspending the drugs in the 1:1:18 solution (5% ethanol, 5% emulphor, and 90% of 0.9% saline) at their respective concentrations for each experiment. The Liposomal blank and liposomal KT109 formulations at desired doses were prepared by diluting the stock formulations in PBS accordingly.

#### 2.5.5 Establishing CIPN in Mice

After acclimation as described above and prior to any treatment and experimentation, baseline mechanical sensitivity thresholds for all mice were assessed using a von Frey assay (see previously described methods). Mice were then given an intraperitoneal (i.p.) injection of paclitaxel (8 mg/kg) or vehicle every other day for a total of four injections. One week after their final respective injection, von Frey thresholds were assessed to confirm that paclitaxel resulted in mechanical hypersensitivity [17–19].

#### 2.5.6 Assessing Free KT109 and Liposomal KT109 *in vivo* Potency

Mechanical hypersensitivity was assessed in all mice 1 hour after each respective dose was i.p. administered for both KT109 and liposomal KT109. Using our methodology for the behavioral assessment of mechanical hypersensitivity, we assessed KT109 (4, 12, or 40mg/kg) in a counter-balanced fashion with 12 mice. Shortly after assessing KT109 at these doses, another experiment used an additional 6 mice to test KT109 (80 mg/kg) and 6 vehicle treated mice. A separate experiment assessed liposomal KT109 (0.04, 0.12, 0.24, or 0.4mg/kg) or liposomal blank in a counter-balanced fashion using the same methods above with 12 mice. One hour after injections, all mice were assessed for mechanical allodynia.

#### 2.5.7 Liposomal KT109 Time Course Evaluation

A semi-Latin square design was employed in which liposomal KT109 (0.4 mg/kg) or liposomal blank was i.p. administered, and mice were given a single von Frey assessment at 0.5, 1, 2, 4, 8, or 24 h post injection. We collected the 48 hours post injection timepoint by i.p. injecting the same group of mice used to assess the previous timepoints with liposomal KT109 (0.4 mg/kg) or liposomal blank which was followed by assessing mechanical hypersensitivity in all mice after 48 hours. A minimum wait time of 72 hours between each treatment was done to prevent carryover effects.

#### 2.5.8 Liposomal KT109 Repeated Dosing Regimen

Mice were i.p. administered liposomal blank or 0.4 mg/kg liposomal KT109 once a day for seven consecutive days. On day 7, all mice were assessed for von Frey thresholds 1 hour after their final injection.

#### 2.5.9 Challenging Liposomal KT109 with Dorsomorphin

Prior to collecting our presented data, all mice had daily i.p. dorsomorphin injections for two consecutive days with dorsomorphin being unsuspended and undissolved in 0.9% saline but this was ended after day two due to concerns with experimental outcomes. Dorsomorphin in a suspension 1:1:18 (v/v/v) solution (200 proof ethanol: emulphor: 0.9% saline) was then used for the following. Mice received a single daily i.p. injection of dorsomorphin (20 mg/kg) or vehicle for five consecutive days. Immediately after the final injection of dorsomorphin or vehicle, mice were given an i.p. injection of liposomal KT109 (0.4 mg/kg) or liposomal blank. Mice were then assessed for mechanical hypersensitivity 1 hour after administration of liposomal KT109 or liposomal blank.

### 2.6 Gel-Based Competitive Activity Based Protein Profiling (ABPP)

To test whether DAGLβ was inhibited in selected tissues, lysates extracted from these tissues were subjected to competitive ABPP with vehicle (DMSO), KT109 (10 µM) and KT195 (10 µM). KT195 is used to account for KT109 off-targets [5]. These compounds were utilized together in the same fashion as previous studies to confirm the presence and activity of DAGLβ in lysate [8]. Brain and spleen tissues from female wildtype mice treated with paclitaxel and liposomal blank, paclitaxel and Liposomal KT109, and untreated female DAGLβ^-/-^mice were used for comparison. For each tissue, insoluble tissue lysates were pre-treated with compound for 30 min at 37 °C, and then treated with HT-01 at 1 µM for 30 min at 37 °C. The reaction was quenched using SDS-PAGE loading buffer, separated via SDS-PAGE (10% acrylamide), and visualized via in-gel fluorescence scanning on a Chemidoc MP imaging system. Protein loading controls were determined through stain free blot scanning on the same system.

### 2.7 Statistical Analysis

All statistical analyses were performed using GraphPad Prism 10.1 (GraphPad Software Inc., San Diego, CA, USA). Student’s t-test was used for comparing two experimental groups, and one-way or two-way ANOVAs were used for comparing three or more groups, followed by Tukey’s post-hoc test where appropriate. Differences were considered statistically different if p < 0.05. For chemoproteomics, protein identification and relative quantification was determined using Proteome Discoverer 3.0 workflow with a PMI-Byonic node as previously described [16,20].

## 3. Results

### 3.1 Second generation liposomal KT109 shows improved drug encapsulation efficiency and stability over time

In this work, we designed an improved, second-generation liposomal KT109 formulation by altering the lipid composition and refining manufacturing methods. The new formulation is composed of DSPC, DSPE-PEG2000, and Cholesterol at molar ratios of 50.7, 7.5, and 41.8%, respectively, versus the DSPC, DSPE-PEG2000, and DOPE which were used in the first-generation formulation (**Fig. 1A**, **1B**). Additional details on the formulation composition can be found in **Table S1 in Supplemental Information**. This composition was adapted from and inspired by other approved liposomal products, using highly compatible lipids to form successful systems [9,10]. EE of KT109 was significantly improved compared to the previous formulation [8], showing nearly 40% EE. Size and spherical morphology of the liposomal KT109 was further confirmed via Cryo-TEM imaging (**Fig. 1C**). Morphology of the liposomal blank formulation was previously reported by the team [11] and is consistent with that of the liposomal KT109.

**Fig. 1.**
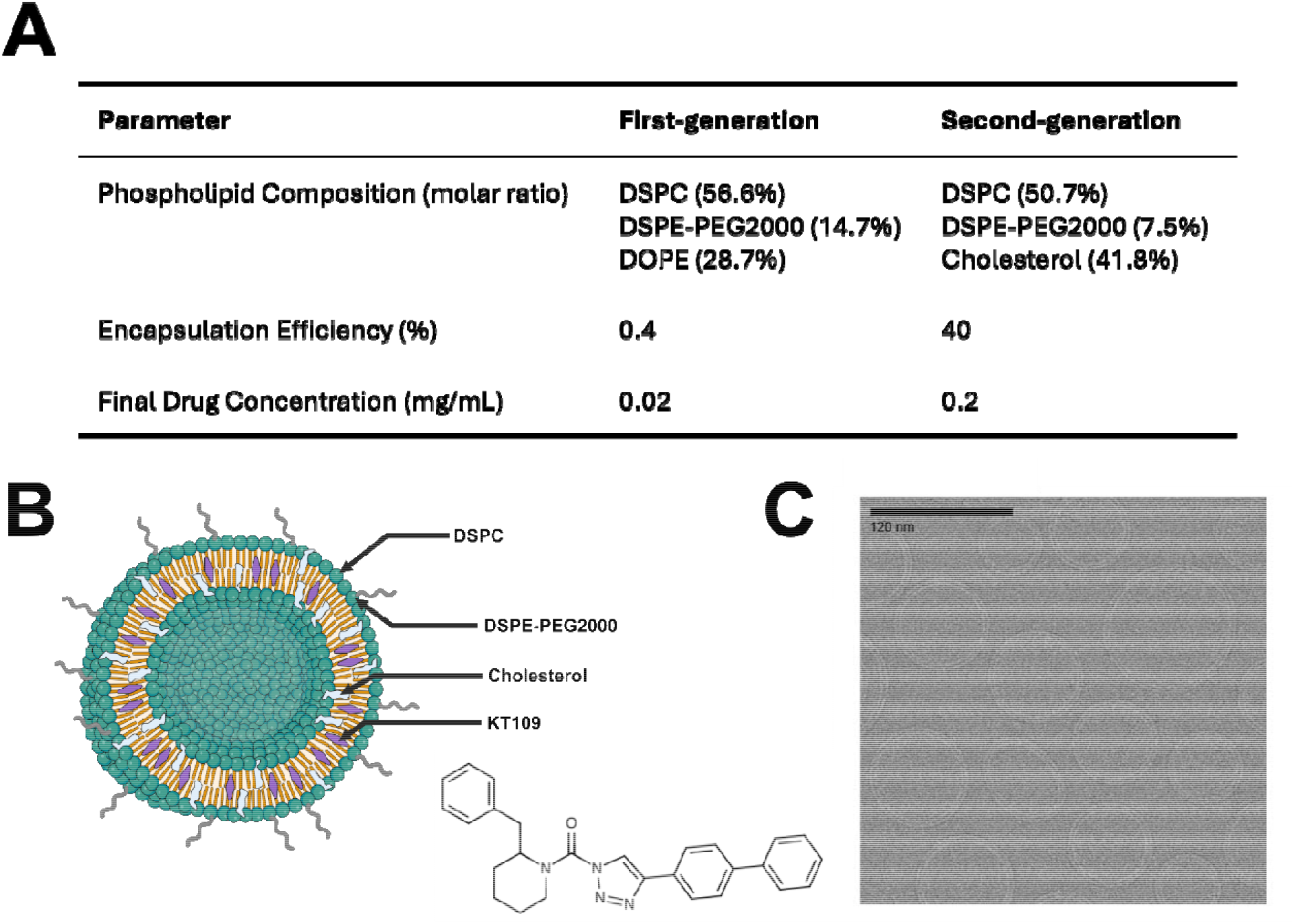
Overview of Improved Liposomal KT109 Formulation. **(A)** Composition and manufacturing efficiency of the improved Liposomal KT109 compared to the first-generation formulation [8] **(B)** Schematic of Liposomal KT109 and drug structure. Hydrophobic KT109 is contained within the phospholipid bilayer, formed by DSPC, DSPE-PEG2000, and Cholesterol. **(C)** Cryo-TEM image of Liposomal KT109. The scale bar represents 120 nm.

Both the second-generation liposomal blank and KT109 were able to achieve average size around 120 nm, PDI < 0.1, and zeta potential near -10mV, indicating stable and uniform liposomal suspensions. Although PDI and zeta potentials were not reported for the first-generation formulation, the droplet size for our new formulation is comparable [8]. A comprehensive stability study was not performed on the first-generation formulation, however, both our second-generation liposomal blank and KT109 formulations showed minimal fluctuations in these key physicochemical properties when stored at 4 for 120 days, indicating good stability over time and drug product feasibility (**Fig. 2A-C**). ZetaView analysis provided additional insight into the systems, further confirming the particle size distributions and concentrations. These results were also used to calculate the lipid concentration (molecules) per liposome (1.36E6 for both), as well as the number of KT109 molecules per liposome (7.6E4). (**Fig. 2D**). Further details surrounding the ZetaView analysis of the liposomal blank was reported in previous work [11]. The second generation liposomal KT109 achieved a much higher concentration (0.2 mg/mL) than the previous first generation liposomal KT109 (only 0.02 mg/mL), which remained stable over the duration of the 120 days (**Fig. 2E**). Notably, the formulation’s key physicochemical characteristics were also comparable across three independent batches (**Table S2 in SI in Supplemental Information**), indicating formulation reproducibility.

**Fig. 2.**
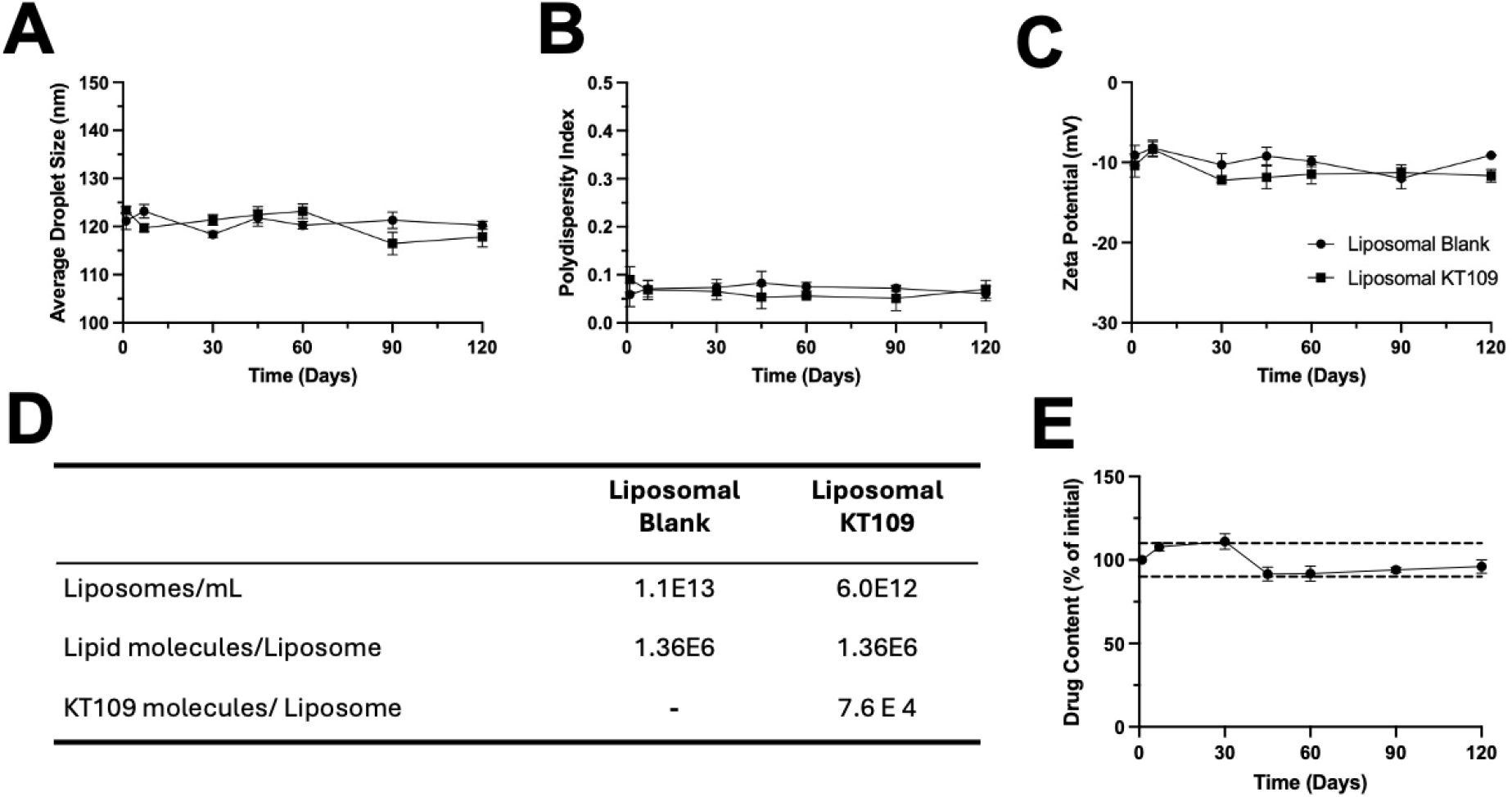
Characterization of Liposomal Blank and KT109 formulations. **(A)** Average droplet size, **(B)** Polydispersity Index, and **(C)** Zeta Potential remain stable upon storage at 4 for up to 120 days, as measured via Zetasizer. Data are presented as Mean ± SD (n=3). **(D)** Key concentrations revealed via ZetaView analysis for both liposomal KT109 and liposomal Blank. [11] **(E)** Drug content of liposomal KT109 remains stable upon storage at 4 for up to 120 days. Data are presented as Mean ± SD (n=3) and are expressed as percent of initial drug content. Black dashed lines indicate 90 to 110% of the initial drug content.

### 3.2 Liposomal KT109 shows reduced potency against known off-targets of KT109 in macrophage cell line, RAW264.7

Previous liposomal KT109 formulations were not tested in a cellular context due to low EE [8]. With newly enhanced EE, we began our investigation of liposomal KT109 treatments in a RAW264.7 macrophage cell line in the context of classical LPS inflammatory stimulation. LC-MS/MS-based competitive activity-based protein profiling (ABPP) evaluation of KT109 versus liposomal KT109 formulation for serine hydrolase inhibition over an extended treatment time (24 hours) demonstrated improved selectivity for DAGLβ (**Fig. 3**). Specifically, known off-targets potently inhibited by free KT109, including platelet-activating factor acetylhydrolase 2 (PAFAH2) and phospholipase A2 Group XV (PLA2G15) [5], were unaffected by liposomal KT109 treatment (**Fig. 3A** and **B**). KT109 showed inhibitory effects on fatty acid amide hydrolase (FAAH) under these chronic treatment conditions, which were reduced with liposomal KT109 (**Fig. 3A**). Taken together, these results demonstrate that the second generation of liposomal KT109 inhibits DAGLβ in macrophages while having reduced potency for KT109 off-targets with extended treatment. With this confirmation of efficacy, we sought to continue this investigation *in vivo* with new behavioral pain models.

**Fig. 3.**
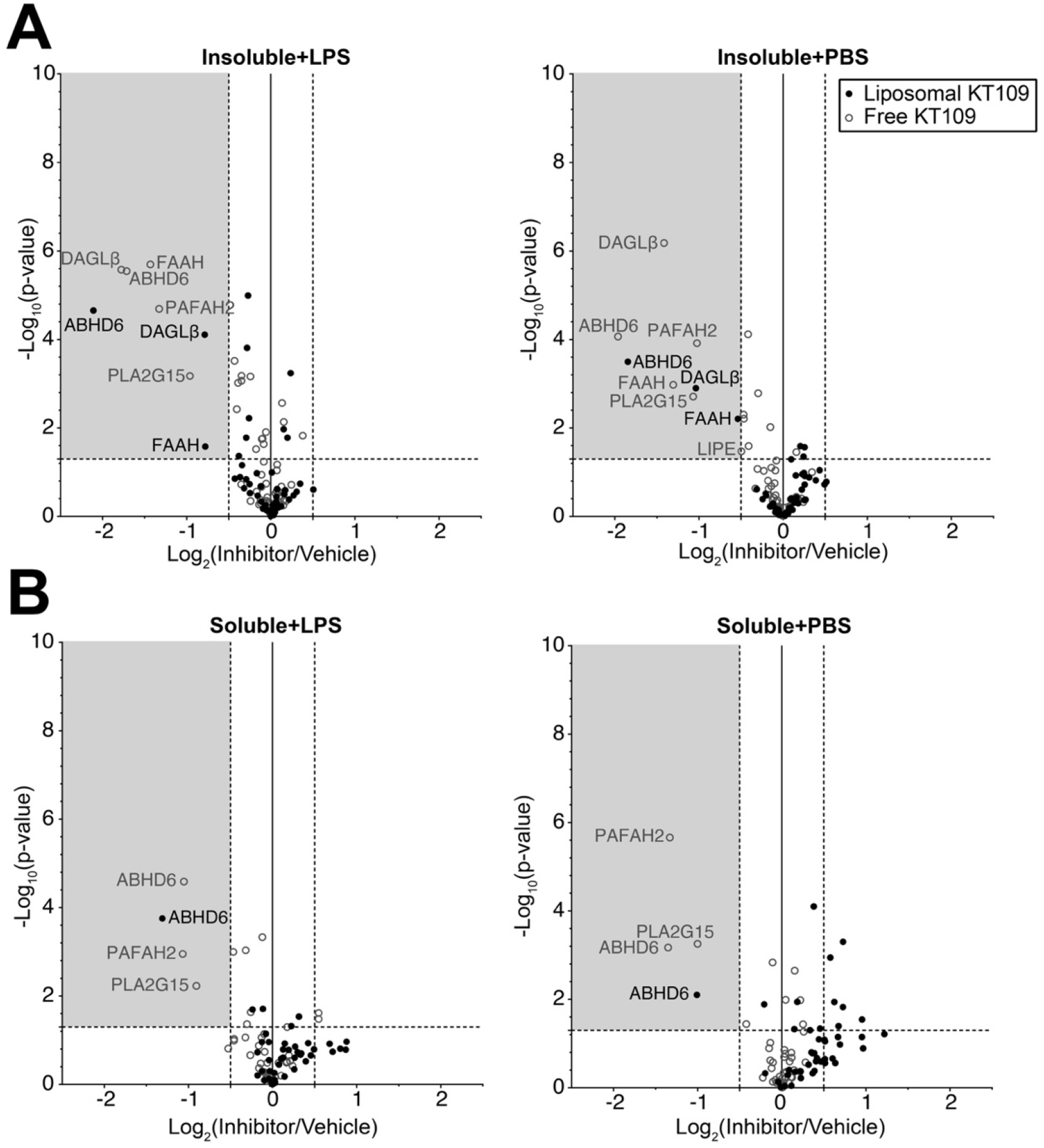
Liposomal KT109 reduces off-target reactivity and potency in murine macrophage cell line, RAW264.7, in both basal and acute inflammatory states. Quantitative activity-based protein profiling of **(A)** insoluble and **(B)** soluble proteomes from inhibitor-vs vehicle-treated cells was performed via TMT-based LC-MS/MS chemoproteomics workflow as described in the Methods section. Data is presented as volcano plots of protein abundance ratios between proteomes from liposomal KT109 (black closed circle, 200 nM) or free KT109 (open circle, 200 nM)-treated over vehicle (PBS)-treated cells. Dashed lines indicated statistical significance cutoff (horizontal: p=0.05) and fold change cutoff (vertical Log_2_FC=±0.5) of inhibitor/vehicle comparisons. Left quadrant (grey box) represents decreased protein abundance in inhibitor-treated samples, n=3 biological replicates in technical duplicate.

### 3.3 Liposomal KT109 shows a dramatic increased antinociceptive potency compared to free KT109 which is not a consequence of off-target delivery

**Fig. 4A** shows the experimental procedure for testing liposomal KT109 and free KT109 in the paclitaxel model of neuropathic pain. Free KT109 (**Fig. 4B**) and liposomal KT109 (**Fig. 4C**) dose-dependently reversed paclitaxel-induced mechanical hypersensitivity. These data were used to calculate an effective dose 50 (ED50) value for free KT109 as well as liposomal KT109 which revealed that liposomal KT109 is several orders of magnitude (164-fold (95% CL: 88-314) more potent than free KT109. The calculated ED50 [95% CL] values were 31 [21–46] mg/kg for free KT109. The calculated ED50 [95% CL] values were 0.226 [0.113-0.454] mg/kg for liposomal KT109. Liposomal KT109 (0.4 mg/kg) fully reversed paclitaxel-induced allodynia at 1 hour post administration with its effects lasting up to 24 hours. In contrast, liposomal blank did not affect von Frey thresholds in paclitaxel treated mice (**Fig. 4D**).

**Fig. 4.**
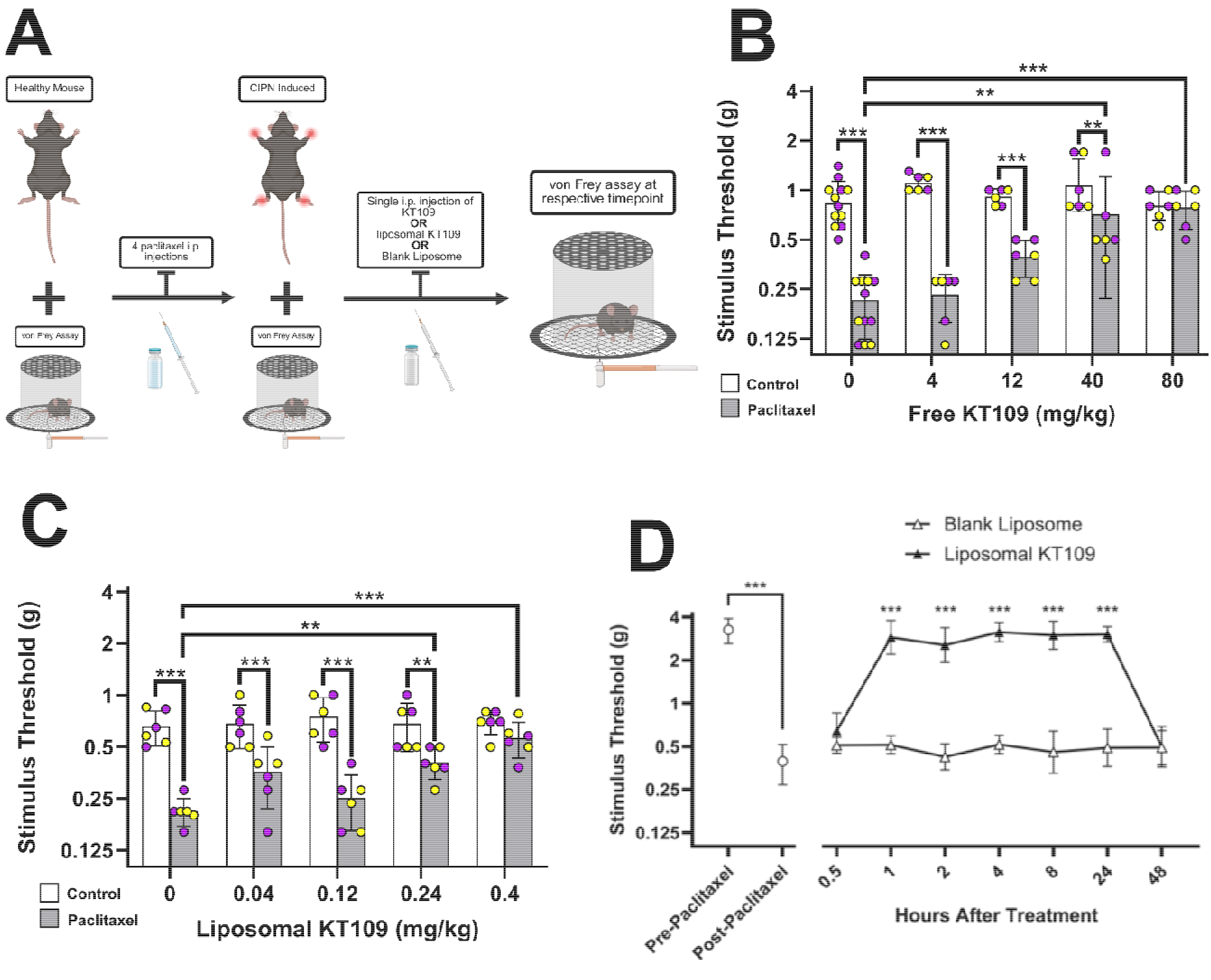
Liposomal KT109 has substantially increased antinociceptive potency compared to free KT109, with antinociceptive effects sustained for 24 hours. **(A)** Schematic overview of CIPN establishment, treatment administration, and behavioral assessment. **(B)** An i.p. injection of free KT109 (80 mg/kg) fully reversed mechanical hypersensitivity induced by paclitaxel 1 hour after administration. Bars show mean ± SD. n = 6-12 mice for vehicle and 6 mice/dose. **(C)** An i.p. injection of liposomal KT109 (0.4 mg/kg) fully reversed the mechanical hypersensitivity produced by paclitaxel 1 hour post administration. Bars show mean ± SD. n = 6-12 mice for vehicle and 6 mice/dose. **(D)** An i.p. injection of liposomal KT109 (0.4 mg/kg) fully reversed the mechanical hypersensitivity 1 hour after administration with this effect lasting for 24 hours. n = 16 for both pre-paclitaxel and post-paclitaxel groups. n = 8 for each liposomal blank and liposomal KT109 point. A two-way ANOVA was utilized to show statistical significance. *** p < 0.001 versus each group’s respective liposomal blank timepoint. For panels B and C, scatter plots reflect data points from individual mice with bars depicting mean ± SD. Yellow circles representing female mice and purple circles reflecting male mice. A Two-way ANOVA was used to assess statistical significance. ** p < 0.01. *** p < 0.001. For the time course study all liposomal KT109 time points were statistically compared to its own liposomal blank timepoint.

To validate that these observed changes in behavior were not a consequence of central nervous system (CNS) activity, we performed ABPP analysis on tissues collected from paclitaxel- and liposomal blank versus liposomal KT109-treated mice. Specifically, we utilized *in vitro* competitive gel-based ABPP with KT109 and its established negative (DAGLβ-inactive) control, KT195.[5] This allowed us to delineate between DAGLβ and an overlapping, unidentified serine hydrolase with the same molecular weight (∼70kDa) that was inhibited by KT109 and KT195 in DAGLβ*^+/+^*and untreated DAGLβ*^-/-^*mice (**Fig. S1, Supplemental Information**). Using this approach, we demonstrated that the DAGLβ band was not affected in liposomal blank or liposomal KT109 treatments, indicating a lack of DAGLβ inhibition in the brains of treated mice (**Fig. S1A, Supplemental Information**). We also confirmed that DAGLβ was not inhibited in immunological organs like the spleen (**Fig. S1B, Supplemental Information**).

### 3.4 Liposomal KT109 sustains its antinociceptive effects after repeated administration

Given that paclitaxel can elicit chronic neuropathic pain in patients, which would require prolonged treatment of an analgesic, we next tested whether repeated administration of liposomal KT109 would maintain its antinociceptive effects of undergo tolerance (**Fig. 5A**). Paclitaxel induced mechanical hypersensitivity was fully reversed with acute i.p. administration of liposomal KT109 compared to its control groups (**Fig. 5B**). Additionally, daily i.p. administration of liposomal KT109 for seven consecutive days did not alter the antinociceptive effects similarly observed in acute dosing (**Fig. 5B)**.

**Fig. 5.**
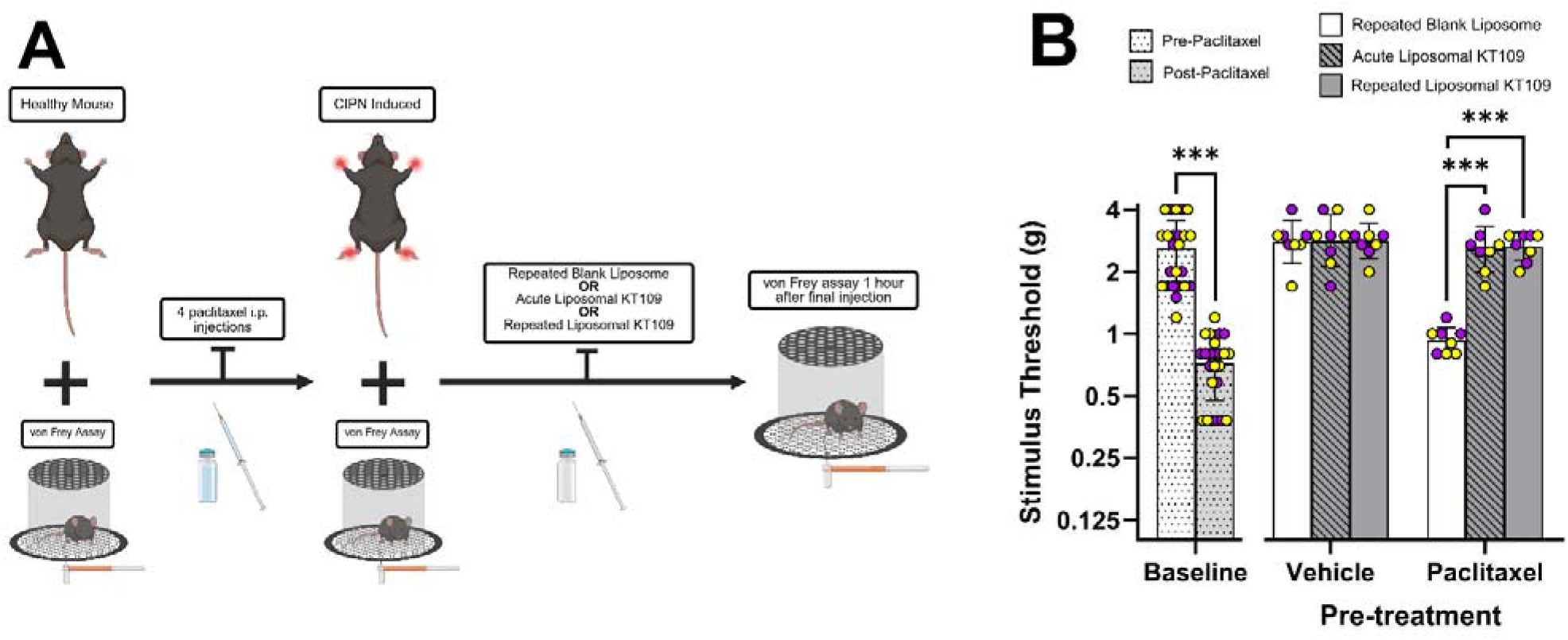
Liposomal KT109 retains its antinociceptive effects with repeated dosing. **(A)** Schematic overview of CIPN establishment, treatment administration, and behavioral assessment. (**B**) A daily i.p. injection of liposomal KT109 (0.4 mg/kg) for seven consecutive days fully maintained its effectiveness in reversing paclitaxel-induced mechanical allodynia. All data are shown as a scatter plot with yellow circles representing female mice and purple circles reflecting male mice. Bars show mean ± SD. n = 48 for both baseline groups. n = 8 per group. Significant Paclitaxel X Treatment interaction (F(2, 42) = 11.10, p = 0.0001). *** p < 0.001.

### 3.5 AMPK plays a role in antinociceptive effects produced by DAGLβ inhibition

We previously reported that the AMPK inhibitor dorsomorphin reversed the antinociceptive phenotype of DAGLβ KO mice in an LPS model of inflammatory, establishing crosstalk between these pathways [12]. Accordingly, we tested whether this regulation extends to the paclitaxel neuropathic pain model **(Fig. 6A)**. As shown in **(Fig. 6B)**, dorsomorphin prevented the antinociceptive effects of liposomal KT109. Additionally, dorsomorphin did not affect von Frey thresholds in mice administered liposomal blank. Thus, with inhibited AMPK preventing antinociception, these data suggest that AMPK mediates antinociception produced by liposomal KT109.

**Fig. 6.**
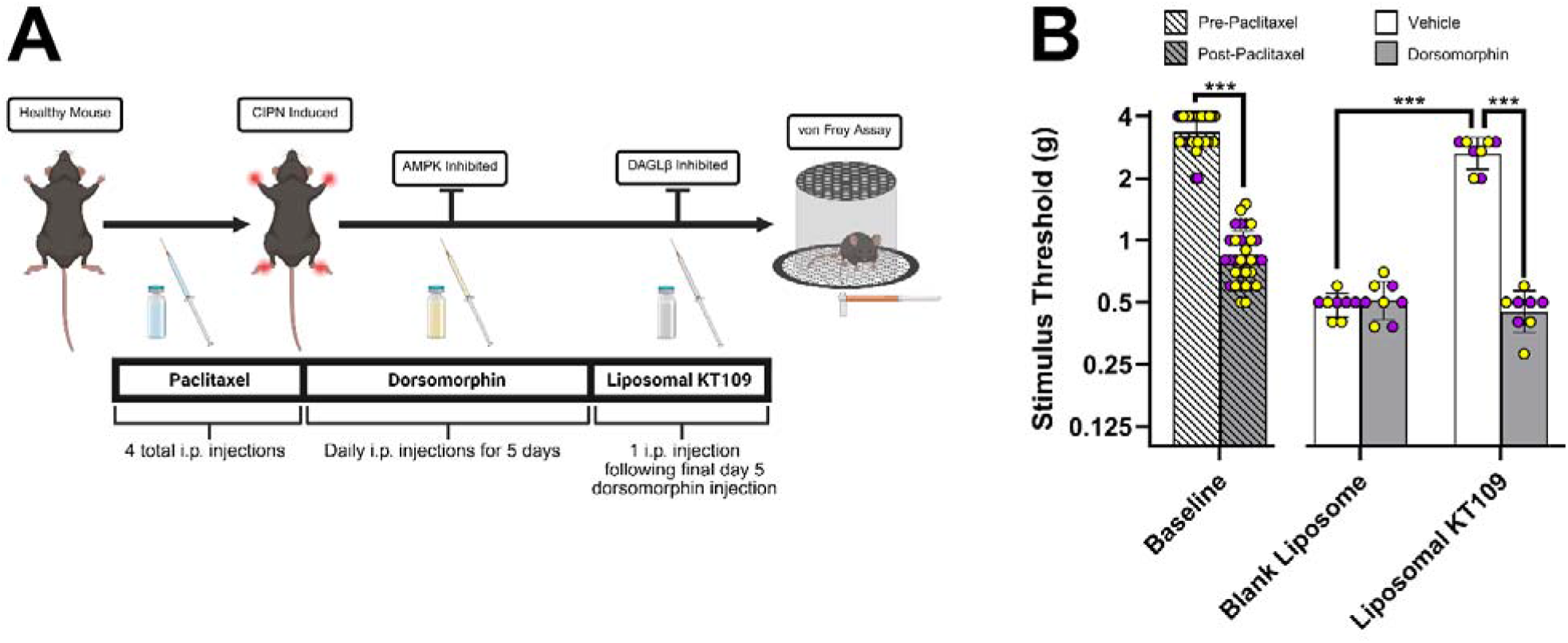
AMPK mediates the antinociceptive effects produced Liposomal KT109. (**A**) Schematic overview of CIPN establishment, treatment administration, and behavioral assessment. (**B**) Dorsomorphin (20 mg/kg) did not alter mechanical hypersensitivity von Frey thresholds in paclitaxel-treated mice prior to liposomal KT109 (0.4 mg/kg) administration. The antinociceptive effects produced by intraperitoneally injected liposomal KT109 (0.4 mg/kg) in paclitaxel-treated mice 1 hour after treatment was inhibited by 5 consecutive days of intraperitoneal dorsomorphin (20 mg/kg) injections prior to liposomal KT109 injections. All data is shown as a scatter plot with yellow circles representing female mice and purple circles reflecting male mice. Bars show mean ± SD. A Two-way ANOVA was utilized to show statistical significance. n = 8 mice/group; *** P < 0.001 vs vehicle and liposomal KT109 groups. No statistical significance was found between any paclitaxel treated mice including post-paclitaxel.

## 4. Discussion

The first generation of liposomal KT109 served as an essential proof-of-concept for *in vivo* efficacy and delivery [8]. However, improvements were necessary for further investigation. Optimization and changes to the liposomal composition led to the creation of a second generation of liposomal KT109 with 100-fold increase in EE compared to first generation counterpart (**Fig. 1**). With improved EE and long-lasting shelf stability (**Fig. 2**), we began by validating that our new liposomal KT109 not only inhibited DAGLβ in macrophages, but did so with reduced off-target reactivity and potency in both basal and acute inflammatory contexts compared to free KT109 (**Fig. 3**). Following this confirmation, we moved into investigating this new and improved formulation in the context of neuropathic pain. Similar to the previous formulation, our new liposomal KT109 showed a striking increase (164-fold) in antinociceptive potency compared to free KT109 (**Fig. 4**). This effect was maintained even after repeated administrations suggesting a lack of tolerance (**Fig. 5**). In alignment with previous studies with free KT109 [8,12], liposomal KT109 antinociception was also reversed with dorsomorphin demonstrating that AMPK-DAGLβ crosstalk and regulation extend to our CIPN mouse model (**Fig. 6**). We further demonstrated that the antinociception we observed in our CIPN mouse model was not a consequence of DAGLβ inhibition within the central nervous system or immunological tissues like the spleen (**Fig. S1, Supplemental Information**).

Our team’s previous work led to the development of the first liposomal KT109 formulation consisting of DOPE, DSPC, and DSPE:PEG2000 [8], which exhibited relatively low drug EE (only 0.4%). Moreover, the first liposomal KT109 did not include cholesterol, the important component to stabilize the liposomal bilayer structure. Although it showed antinociceptive potency in an inflammatory mouse model and nearly 80% inactivation of DAGLβ in macrophages supporting its therapeutic potential, the practical limitations of the first generation liposomal KT109 formulation stood to be further improved in terms of drug EE, stability and reproducibility. In this study, we successfully developed a new and improved liposomal KT109 formulation by using and adjusting the relevant lipid compositions from FDA-approved liposomal products such as Onivyde [10] and Doxil [9]. DSPC, DSPE:PEG2000, and cholesterol were selected for their ability to work in tandem to support the bilayer structure and performance. More specifically, DSPC serves as the structural backbone, providing a rigid bilayer through its saturated acyl chains and high transition temperature. Cholesterol sits between DSPC to condense the membrane, facilitate packing, ultimately ensuring cargo retention as well as overall membrane stability. Finally, DSPE:PEG2000 equips the liposome with a hydrated PEG corona that helps to prevent particle aggregation and potentially prolong the circulation of liposomes in the bloodstream. As a testament to this improved composition, our new liposomal KT109 exhibited excellent reproducibility, homogeneity, and stability up to 120 days, supporting its feasibility as a potential drug product (**Fig. 2**) [15]. Manufacturing improvements compared to the first-generation formulation [8] should be noted as well, leading to a 100-fold increase in EE (40% for this formulation compared to 0.4% for the first-generation formulation) as well as an increase in KT109 concentration achieved over 10-fold (0.2 mg/mL versus 0.02 mg/mL; **Fig. 1**). Some of the improvements that were made include adapting the solvent system (chloroform:methanol 1:1 vs chloroform only), increasing formulation volumes (20mL vs <5mL total working volumes), and using a more translatable extrusion system (Genizer GJE-10mL Jacketed Extruder vs 1mL handheld extrusion syringe).

As a consequence of increased EE of second generation liposomal KT109, we were able to test the new liposomal KT109 alongside liposomal blank formulation in live cells before moving to an *in vivo* model [11]. Here, we discovered that with extended treatment, equal concentrations of encapsulated KT109 showed a heightened selectivity for DAGLβ in basal and inflamed state murine macrophages confirming liposomal KT109 functions as a DAGLβ inhibitor within macrophage cells with notably benefits outside of lower effective dosage effects (**Fig. 3**). At the same time, previous research into the liposomal blank formulation revealed a non-zero immunomodulatory and enzymatic effect [11] which emphasized the need for liposomal blank to be used as a liposomal KT109 vehicle control within *in vivo* investigations. Moreover, the absence of detectable DAGLβ inhibition in whole-brain and spleen **(Fig. S1, Supplemental Information)** suggests that these tissues may not have contributed to the antinociceptive effects of liposomal KT109 in our mouse model of CIPN. However, the use of whole-organ homogenates may have obscured our ability to detect DAGLβ inhibition in specific cell populations, such as microglia and macrophages.

When explored *in vivo*, our results revealed that 1 hour after an i.p. injection of liposomal KT109 mechanical hypersensitivity was fully reversed in a dose response fashion with 0.04 and 0.12 mg/kg failing to reverse mechanical hypersensitivity, 0.24 mg/kg partially reversing mechanical hypersensitivity, and 0.4 mg/kg of liposomal KT109 fully reversed mechanical hypersensitivity. A similar trend was observed 1 hour after an i.p. injection of free KT109 where 4, 12, and 24 mg/kg did not reverse mechanical hypersensitivity, 40 mg/kg partially reversed mechanical hypersensitivity, and 80 mg/kg fully reversed mechanical hypersensitivity. Doses for free and liposomal KT109 that fully reversed mechanical hypersensitivity were selected for subsequent studies. Additionally, utilizing these data, we calculated the potency of liposomal KT109 to be 164 times more potent than free KT109 (**Fig. 4**). This impressive *in vivo* potency increase is similar to previously published findings where a differently formulated liposomal KT109 fully reversed mechanical hypersensitivity in an LPS mouse model, however, a dose in mg/kg was not confirmed [8] Notably, previous studies suggest that the marked potency increase of liposomal KT109 could be attributed to the uptake of liposomal KT109 by tissue resident and recruited macrophages which were found to have high expression of DAGLβ [5,8].

Subsequent *in vivo* investigations we conducted showed that liposomal KT109 at a dose of 0.4 mg/kg fully reverses mechanical hypersensitivity 1 hour after being i.p. administered with these effects lasting for 24 hours (**Fig. 4**). Further experiments we performed confirmed that liposomal KT109 sustains its antinociceptive effects following repeated i.p. administration (**Fig. 5**). This finding is similar to previously published results where free KT109 sustained its antinociceptive effects after repeated administration in an LPS mouse model [6]. Together, our liposomal KT109 time course and repeated administration studies indicate a reduced need for frequent dosing with a lack of tolerance development.

As seen in the *in vivo* tissue analyses from mice treated with the first-generation liposomal KT109 formulation [8], the new liposomal KT109 showed a lack of DAGLβ inhibition in spleen and brain (**Fig. S1**). This suggests that our new liposomal KT109 does not confer antinociception through the CNS nor immunological organs like the spleen. We have previously established that DAGLβ-AMPK crosstalk is essential for paclitaxel-induced dorsal root ganglia (DRG) neuronal hyperexcitability [12]. While macrophage infiltration and accumulation within the DRG is known to contribute to the initiation and continuation of paclitaxel-induced neuropathy and neuronal hyperexcitability [21,22], the CIPN model and DRG tissues, unlike previous acute inflammatory models used to study liposomal KT109 [8], do not elicit a highly concentrated population of immune cells (i.e. peritoneal macrophages). Future studies, wherein a larger sample size of mice is utilized and DRG macrophage isolation and sorting procedures are incorporated [22,23], could allow for a concentrated enough population of DRG macrophages to quantitate the activity and expression through LC-MS/MS chemoproteomics and proteomics [16].

Previous mechanistic electrophysiological studies demonstrated a significant increase in DRG neuronal excitability in paclitaxel-treated DAGLβ WT mice compared to untreated DAGLβ WT mice, untreated DAGLβ KO mice, and intriguingly paclitaxel-treated DAGLβ KO mice [12]. More interestingly, parallel electrophysiological studies revealed that paclitaxel-treated DAGLβ KO mice administered dorsomorphin, an AMPK inhibitor, exhibited significantly greater DRG neuronal excitability than paclitaxel-treated DAGLβ KO mice administered vehicle [12]. Additionally, LPS-treated DAGLβ KO mice which were administered dorsomorphin led to a reversal of the innate antinociceptive phenotype observed in DAGLβ knockout mice [12]. Collectively, these previously published mechanistic findings set up the rationale for us to study the antinociceptive role of AMPK following liposomal KT109 treatment in a CIPN mouse model. Here, we found that the inhibition of AMPK blocks the antinociceptive effects produced by liposomal KT109 in a mouse model of CIPN (**Fig. 6**). Although the in-depth mechanism of how AMPK mediates the antinociceptive effects following DAGLβ inhibition remain unclear, our findings nicely complement the previous electrophysiological studies and further suggest that AMPK signaling is necessary for the antinociceptive effects produced by DAGLβ inhibition in neuropathic pain.

Consistent with findings obtained with the first-generation liposomal KT109 formulation [8], the new formulation produced no detectable DAGLβ inhibition in spleen or whole brain (**Fig. S1, Supplemental Information**). These findings suggest that the antinociceptive effects of liposomal KT109 are not mediated through direct action in these organs, though the possibility of cell-specific effects (e.g., microglia and macrophages) cannot be excluded. In CIPN mice, we demonstrated a 164 times greater potency with liposomal KT109 compared to free KT109 without observing tolerance issues. Moreover, we established that this antinociceptive effect is mediated through the activation of AMPK as seen in other pain models with free KT109 [12]. While further research into the site and mode of action of this potent antinociception are critical and represent an ongoing research effort, these findings represent a foundation and working model for continued exploration of liposomal KT109 in other chronic pain models and of other liposomal-encapsulated small molecules for *in vivo* delivery.

## 5. Conclusions and Future Directions

In this work, we report the development and in vivo characterization of a second-generation liposomal KT109 formulation. Adjustments to lipid composition and processing increased EE 100-fold and raised the drug concentration more than 10-fold relative to our first-generation formulation. Activity-based protein profiling showed that the formulation retained DAGLβ potency in both basal and acute inflammatory states while reducing off-target reactivity, confirming improved selectivity for DAGLβ. In a mouse model of neuropathic pain, the formulation produced greater antinociception than free KT109 at equivalent doses, with effects sustained for up to 24 hours. Antinociception was not present when AMPK was inhibited by dorsomorphin, a result consistent with an AMPK-dependent mechanism of action. Postmortem ABPP analysis also showed that DAGLβ was not inhibited in whole brain or spleen. Although these findings support liposomal KT109 as a promising treatment for neuropathic pain, further work is needed to elucidate the pathway linking DAGLβ inhibition to AMPK-dependent antinociception as well as to confirm translational potential through biodistribution, pharmacokinetic, and chronic dosing studies.

## Supporting information

Supplemental Data Set

## Abbreviations

2-AG: 2-arachidonylglycerol
ABPP: activity-based protein profiling
AMPK: adenosine monophosphate-activated protein kinase
CIPN: chemotherapy-induced peripheral neuropathy
DAGLβ: diacylglycerol lipase-beta
DOPE: 1,2-dioleoyl-sn-glycero-3-phosphoethanolamine
DRG: dorsal root ganglia
DSPC: 1,2-distearoyl-sn-glycero-3-phosphocholine
DSPE-PEG2000: 1,2-distearoyl-sn-glycero-3-phosphoethanolamine-N-[methoxy(polyethylene glycol-2000]
ED_50_: effective dose 50
EE: encapsulation efficiency
FAAH: fatty acid amide hydrolase
FBS: fetal bovine serum
gf: grams of force
HPLC: high-performance liquid chromatography
i.p.: intraperitoneal
LC-MS/MS: liquid chromatography-mass spectrometry/mass spectrometry
LPS: lipopolysaccharide
LUVs: large lamellar vesicles
MLVs: multi-lamellar lipid vesicles
NTA: nanoparticle tracking analysis
PAFAH2: platelet-activating factor acetylhydrolase 2
PBS: phosphate buffered saline
PDI: polydispersity index
PLA2G15: phospholipase A2 Group XV

## Statements and Declarations Acknowledgments

We would like to thank the Sauer Structural Biology Laboratory at The University of Texas at Austin for their help in acquiring cryo-TEM images.

## Funding

This work was supported by National Institute of Health grant (R01DA043571), as well as by the National Institutes of Health grants (DA043571, AI169412, F31NS149625 and P30DA033934), University of Texas at Austin startup funds, and a Research Grant Award from The Welch Foundation (F-2143-20230405 to K.-L.H.).

## Competing Interests

Q.X., A. L., L.B., M.G., R.B., K. H., P.L., are co-inventors on an VCU invention disclosure “DAGLβ inhibitor-loaded liposomes for treating neuropathic pain: compositions and methods of use thereof”, which was filed on 10/15/2026.

## Author Contributions

**Liam A. Babcock, Madeline Gunawardena, Payton E. Lowrey:** data curation, investigation, methodology, formal analysis, writing-original draft. **Rudra Pangeni, McKenzie Hall:** data curation, investigation, methodology, formal analysis, Writing - Review & Editing. **Michael Taylor, Saniya Chethan, Sara M. Herz, Dayna A. Adeso, Olivia L. Murtagh:** data curation, investigation, methodology. **Qingguo Xu, Ku-Lung Hsu, Aron H. Lichtman:** conceptualization, investigation, methodology, writing-review and editing, Funding acquisition.

## Data Availability

All data generated in this work and needed to evaluate the conclusion are presented in the main body of the paper or have been made available in the supplemental information. Additional data that support the findings of this study are available from the corresponding authors upon reasonable request.

## Ethics Approval

All studies involving the use of animals were conducted in compliance with the National Institutes of Health Guide for the Care and Use of Laboratory Animals and approved by the Institutional Animal Care and Use Committee at Virginia Commonwealth University.

