## Supplemental Information for "Liposomal DAGLβ Inhibitor Shows Substantial Antinociceptive Potency Mediated by AMPK in a Neuropathic Pain Mouse Models"

### Table of Contents:

1. Supplemental Tables and Figures
2. Supplemental Dataset 1: Chemoproteomics Log<sub>2</sub>(Fold Change) and p-values (presented in separate document)

**Table S1.** *Liposomal KT109 Composition Details*

The lipid composition utilized for the second generation liposomal KT109 formulation was adapted from those of approved drug products as described in the main text. Table S1 further details each lipid used, along with molar ratios, molecular weights, and masses used for a given 20 mL batch of formulation.

| **Lipid** | **Molar Ratio (%)** | **Molecular Weight (g/mol)** | **Mass (mg)** |
| --- | --- | --- | --- |
| DSPC | 50.7 | 790.2 | 400 |
| DSPE-PEG2000 | 7.5 | 2805.5 | 210.6 |
| Cholesterol | 41.8 | 386.6 | 161.2 |

**Table S2. Liposomal KT109 Batch to Batch Reproducibility**

Liposomal drug products are considered to be reproducible when their properties remain consistent across three independent batches and satisfy predefined quality targets.

Physicochemical properties measured for three separate batches of both the liposomal blank and liposomal KT109 formulations are presented here, alongside the specified quality targets, demonstrating that the formulation meets the criteria for reproducibility.

| *Liposomal Quality Attributes* |  |  |  |  |  |
| --- | --- | --- | --- | --- | --- |
| **Batch** | **Formulation Type** | **Average Droplet Size (nm)** | **PDI** | **Zeta Potential (mV)** | **Encapsulation Efficiency (%)** |
| 1 | Blank | 121.2 ± 1.8 | 0.06 ± 0.03 | -9.1 ± 1.2 | - |
|  | KT109 | 123.4 ± 0.7 | 0.09 ± 0.03 | -10.4 ± 1.4 | 41.7 ± 0.3 |
| 2 | Blank | 119.1 ± 0.5 | 0.05 ± 0.04 | -12.4 ± 1.2 | - |
|  | KT109 | 116.5 ± 2.4 | 0.05 ± 0.03 | -11.3 ± 0.9 | 44.1 ± 0.3 |
| 3 | Blank | 130.3 ± 1.8 | 0.08 ± 0.01 | -10.5 ± 0.2 | - |
|  | KT109 | 131.8 ± 0.9 | 0.07 ± 0.04 | -9.9 ± 0.8 | 45.3 ± 1.8 |
| **Quality Target** |  | **< 200 nm<sup>#</sup>** | **< 0.2<sup>#</sup>** | **-30 to +30 mV<sup>#</sup>** | **> 0.4 %<sup>\*</sup>** |

<sup>#</sup>based on standard liposomal quality targets defined in literature

<sup>\*</sup>based on expected improvement from first generation formulation

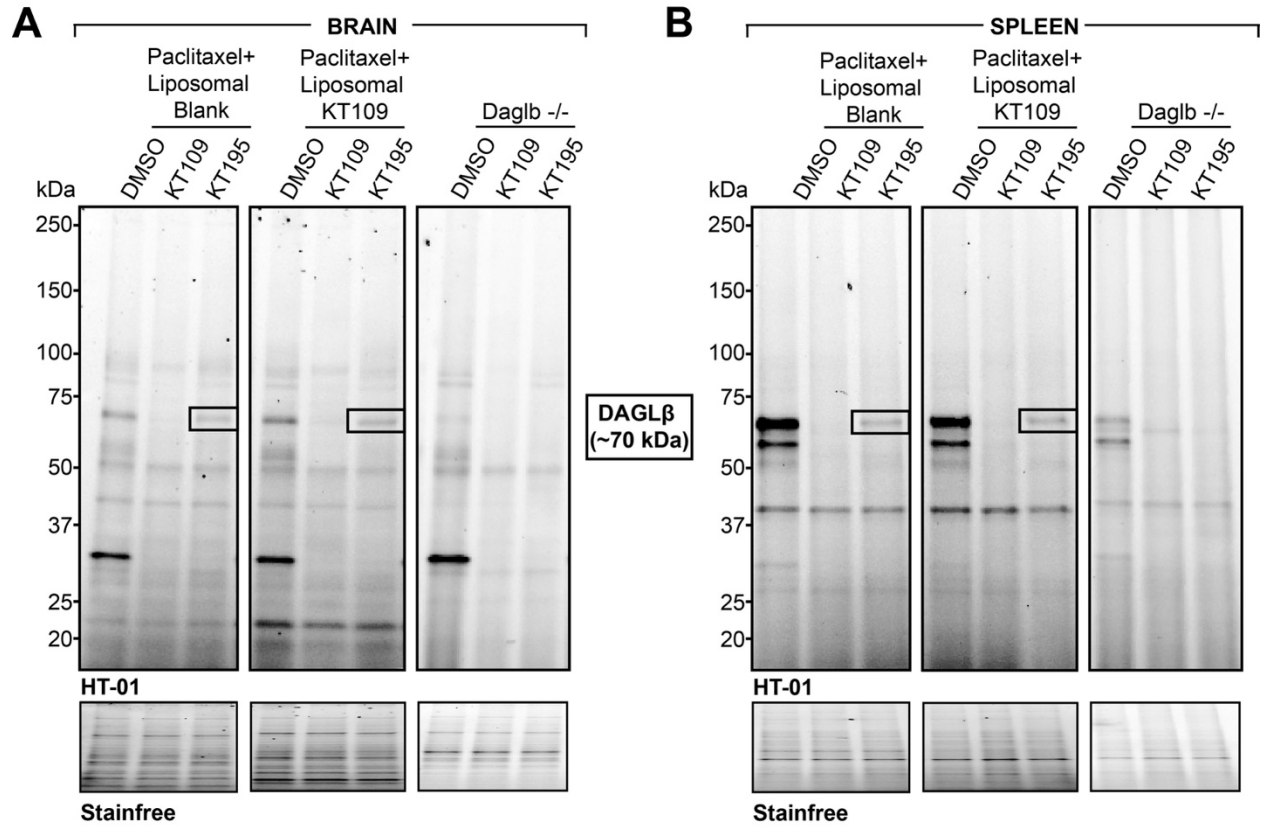

**Figure S1. DAGLβ Identification through *in vitro* competitive ABPP with KT109 and KT195** (A) Brain and (B) Spleen tissue from paclitaxel-treated and liposomal blank or liposomal KT109-treated mice alongside DAGL$\beta^{-/-}$ mice was treated with vehicle (DMSO), KT109 (10 μM), KT195 (10 μM) directly prior to treatment with activity-based probe, HT-01 (1 μM, 30 min, 37°C). DAGLβ band (black box) identified in KT195-treated condition. Stain free represents loading control.
